# Protoporphyrin IX and verteporfin prevent SARS-CoV-2 infection *in vitro* and in a mouse model expressing human ACE2

**DOI:** 10.1101/2020.04.30.071290

**Authors:** Chenjian Gu, Yang Wu, Huimin Guo, Yuanfei Zhu, Wei Xu, Yuyan Wang, Yu Zhou, Zhiping Sun, Xia Cai, Yutang Li, Jing Liu, Zhong Huang, Zhenghong Yuan, Rong Zhang, Qiang Deng, Di Qu, Youhua Xie

**Author notes:** These authors contribute equally (Chenjian Gu, Yang Wu, Huimin Guo, Yuanfei Zhu). Corresponding authors, Youhua Xie, Key Laboratory of Medical Molecular Virology (MOE/NHC/CAMS), School of Basic Medical Sciences, Shanghai Medical College, Fudan University, e-mail,; Di Qu, BSL-3 laboratory of Fudan University, School of Basic Medical Sciences, Shanghai Medical College, Fudan University. e-mail,; Qiang Deng, Key Laboratory of Medical Molecular Virology (MOE/NHC/CAMS), School of Basic Medical Sciences, Shanghai Medical College, Fudan University, e-mail,.

## Abstract

The SARS-CoV-2 infection is spreading rapidly worldwide. Efficacious antiviral therapeutics against SARS-CoV-2 is urgently needed. Here, we discovered that protoporphyrin IX (PpIX) and verteporfin, two FDA-approved drugs, completely inhibited the cytopathic effect produced by SARS-CoV-2 infection at 1.25 μM and 0.31 μM respectively, and their EC50 values of reduction of viral RNA were at nanomolar concentrations. The selectivity indices of PpIX and verteporfin were 952.74 and 368.93, respectively, suggesting broad margin of safety. Importantly, PpIX and verteporfin prevented SARS-CoV-2 infection in mice adenovirally transduced with human ACE2. The compounds, sharing a porphyrin ring structure, were shown to bind viral receptor ACE2 and interfere with the interaction between ACE2 and the receptor-binding domain of viral S protein. Our study suggests that PpIX and verteporfin are potent antiviral agents against SARS-CoV-2 infection and sheds new light on developing novel chemoprophylaxis and chemotherapy against SARS-CoV-2.

## Main Text

The infection of SARS-CoV-2 has spread around the world since December 2019. As of July 6, 2020, there are nearly 11 million confirmed cases globally, of which more than five hundred thousand died (https://www.who.int/emergencies/diseases/novel-coronavirus-2019). Although the pandemic has been contained in some countries, the numbers of confirmed cases and deaths worldwide are expected to continue to rise.

SARS-CoV-2 is transmitted through respiratory droplets and close contact, which causes mainly upper and lower respiratory diseases. The majority of infected healthy adults and children only show mild symptoms including cough, fever, fatigue and diarrhea but the elderly with various chronic diseases are at high risk of development of serious diseases including pneumonia, acute respiratory distress, multiple organ failure and shock. At present, the treatment of Coronavirus Disease 2019 (COVID-19) is mostly supportive, including non-specific antivirals and symptom-alleviating therapies. Ventilations and intensive care are required for severe cases, calling for early intervention to prevent symptoms from deteriorating.

*In vitro* experiment showed that remdesivir targeting viral RNA-dependent RNA polymerase (RdRp) effectively inhibited SARS-CoV-2 replication^1,2^. The compassionate use of remdesivir for patients with severe COVID-19 indicated that clinical improvement was observed in 36 of 53 patients (68%)^3^. Remdesivir was reported to shorten the time to recovery in adults hospitalized with COVID-19 and evidence of lower respiratory tract infection in a double-blind, randomized, placebo-controlled trial, though conflicting trial results have also been reported^4,5^. Several repurposed drugs have been tested *in vitro* for inhibition of SARS-CoV-2 infection and some of them were tested in clinical trial^6–9^. Among them, chloroquine and hydroxychloroquine have been shown to inhibit SARS-CoV-2 infection *in vitro*, while the clinical trials of hydroxychloroquine reported controversial results^2,10–12^ The effective concentrations (presented as the concentration for 50% of maximal effect (EC50) on the reduction of viral RNA) of most previously selected drugs are in the micromolar (μM) concentration range. On the other hand, neutralizing antibodies against SARS-CoV-2 are also being intensively studied^13–15^. In general, more efficacious antiviral therapeutic agents against SARS-CoV-2 with good safety profile are urgently needed.

In search of novel antivirals that can effectively inhibit SARS-CoV-2 infection, we set out to screen an FDA-approved drug library of 3200 small molecules via observation of viral CPE in Vero-E6 cells, followed by evaluation of the antiviral effect of candidate compounds in *vitro* and in mice transduced intranasally with the recombinant adenovirus 5 expressing human ACE2 (Ad5-hACE2). We discovered that protoporphyrin IX (PpIX) and verteporfin displayed a potent antiviral activity and prevent SARS-CoV-2 infection.

## Results

### Protoporphyrin IX and verteporfin effectively inhibit SARS-CoV-2 infection in Vero-E6 cells

Two compounds, protoporphyrin IX and verteporfin, showed a complete suppression of viral CPE at 1.25 μM and 0.31 μM respectively (Fig. 1B). These two compounds were subject to further analysis. At 48 hours post-infection, viral RNA level in the supernatant of the compound-treated cells was measured using qRT-PCR, which decreased dose-dependently as the compound concentration increased. Based on the RNA level-compound concentration curve, the EC50 values of protoporphyrin IX, verteporfin and the positive control remdesivir were calculated to be 0.23 μM, 0.03 μM, and 1.35 μM (Fig. 1A), respectively. The EC50 of remdesivir was comparable to the previous report^2^. Cell viability assay was performed, resulting in a viability-compound concentration curve (Fig. 1A), from which the CC50 (cytotoxicity concentration 50%) values of protoporphyrin IX, verteporfin and remdesivir were determined to be 219.13 μM, 10.33 μM, and 303.23 μM, respectively. The selectivity indices (S.I.) for the three compounds could thus be calculated as 952.74, 368.93, and 224.61, respectively. Viral N protein expression in infected Vero-E6 cells was assessed by immunofluorescence. The data revealed the complete inhibition of N protein expression by protoporphyrin IX, verteporfin and remdesivir at 1.25 μM, 0.31 μM, and 6.25 μM, respectively (Fig. 1B). The results indicate that protoporphyrin IX and verteporfin strongly inhibit the infection of SARS-CoV-2 at nanomolar concentrations and have a wide safety range *in vitro*.

**Fig. 1.**
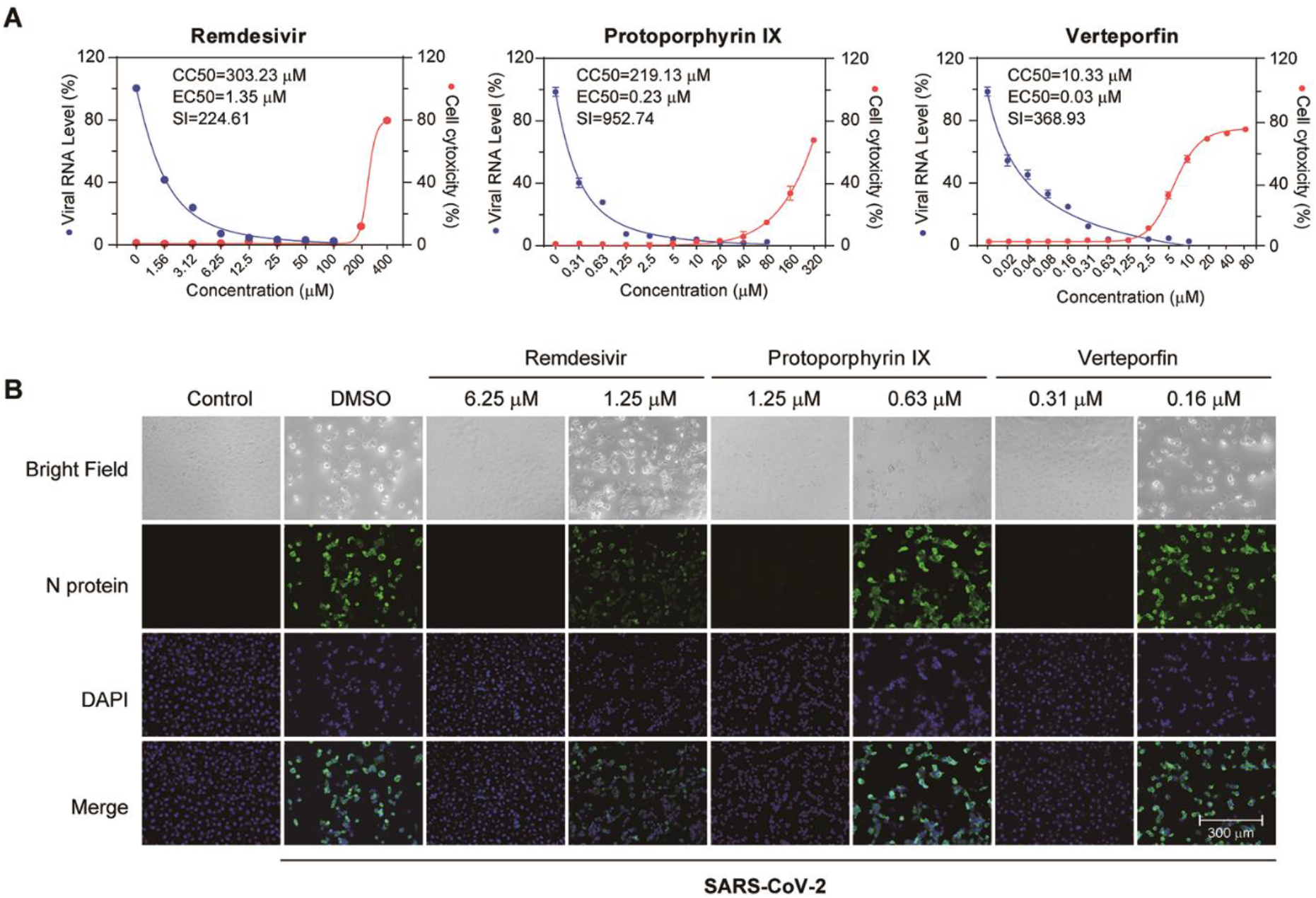
Effective inhibition of SARS-CoV-2 infection by protoporphyrin IX and verteporfin. **(A)** Antiviral effect and cell cytotoxicity of protoporphyrin IX and verteporfin. The viral RNA production in the supernatant of infected Vero-E6 cells was quantified with qRT-PCR. The value at each compound concentration was presented relative to that at zero compound concentration that was set as 100% (blue). The percentage of reduction in viable cells at different compound concentration (red) was measured using the CCK8 assay. The value at each compound concentration was calculated using the formula, 100-Value (compound concentration)/Value (zero compound concentration). EC50, concentration for 50% of maximal effect; CC50, concentration for 50% of maximal cytotoxic effect; S.I., selectivity index. Data from three independent experiments were analyzed. **(B)** Immunofluorescence of intracellular viral N protein. Intracellular expression of N protein was assessed by staining of infected Vero-E6 cells with the polyclonal anti-N antibody (1:1000 dilution, green). Nuclei were stained with DAPI. CPE was shown in bright field.

### Effects of treatment timing on protoporphyrin IX and verteporfin’s inhibition of SARS-CoV-2 infection

We next analyzed the relationship between the antiviral effect and treatment timing of protoporphyrin IX and verteporfin. As shown in Fig. 2A, Vero-E6 cells were treated with protoporphyrin IX, verteporfin or the solvent DMSO before viral infection, during viral entry and after viral entry. A total of 8 treatment groups were set up for each compound (group I-VIII). Based on the previous results, we selected the compound concentrations of 2.5 μM and 1.25 μM for protoporphyrin IX and verteporfin, respectively. At 48 hours post infection, viral RNA level in the culture supernatant was quantified with qRT-PCR. The results showed that viral RNA levels of all the compound-treated groups (group I-VII of each compound in Fig. 2B, 2C) were significantly lower than that of the DMSO-treated group (group VIII in Fig. 2B, 2C). Importantly, pre-treatment alone resulted in the complete inhibition of SARS-CoV-2 infection (group IV in Fig. 2B, 2C). In addition, treatment of cells with protoporphyrin IX or verteporfin after viral infection also inhibited viral RNA production, albeit to different extent (group VII in Fig. 2B, 2C). The results of immunofluorescence analysis on intracellular viral N protein were consistent with those of viral RNA measurement (Fig. 2D). Collectively, the results indicate that protoporphyrin IX and verteporfin can prevent SARS-CoV-2 infection and might suppress established SARS-CoV-2 infection to some degree.

**Fig. 2.**
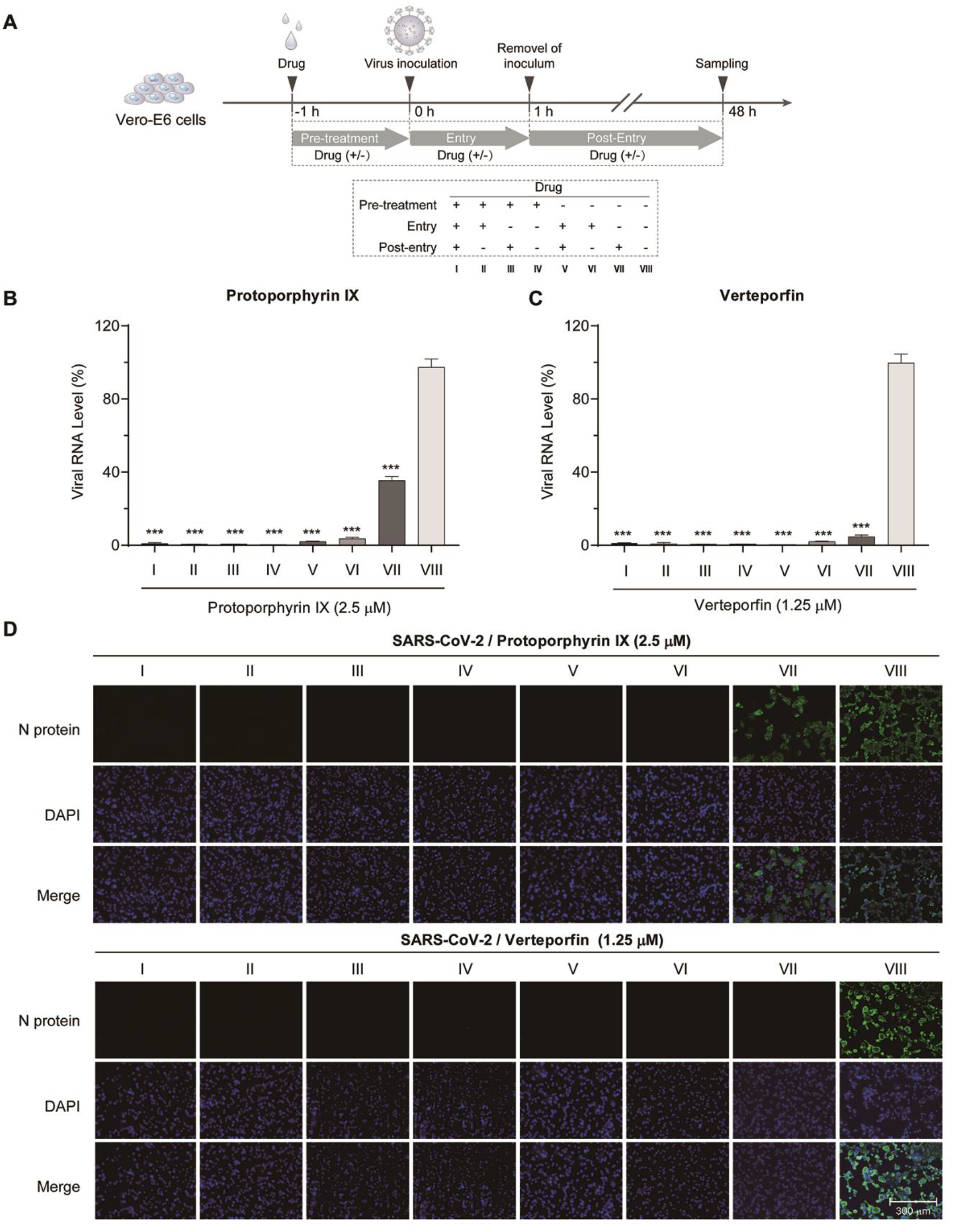
Effects of treatment timing of protoporphyrin IX and verteporfin on SARS-CoV-2 infection. **(A)** Schematic presentation of treatment timing of protoporphyrin IX and verteporfin. Briefly, Vero-E6 cells were treated with protoporphyrin IX, verteporfin or the solvent DMSO before viral infection, during viral entry and after viral entry. A total of 8 treatment groups (I-VIII) for each compound were set up. **(B)** Antiviral effect of different treatment timing. Viral RNA level in the supernatant of infected Vero-E6 cells was quantified with qRT-PCR. The values of group I to VII were presented relative to that of group VIII which was set as 100%, respectively. Statistical significance was determined using the unpaired two-tailed Student’s *t* test. *** *P* < 0.001. Data from three independent experiments were analyzed. **(C)** Immunofluorescence of intracellular viral N protein. Intracellular expression of N protein of different treatment timing was assessed by staining of infected Vero-E6 cells with the polyclonal anti-N antibody (1:1000 dilution, green). Nuclei were stained with DAPI.

The preventive effect was further tested by the pre-treatment of cells with either compound at a constant concentration and later infection with an increasing virus titer (Fig. 3A). As shown in Fig. 3B, no viral N protein expression was detected in protoporphyrin IX or verteporfin pre-treated cells even if the inoculated viral titer was raised by 16 folds (200 PFU to 3200 PFU).

**Fig. 3.**
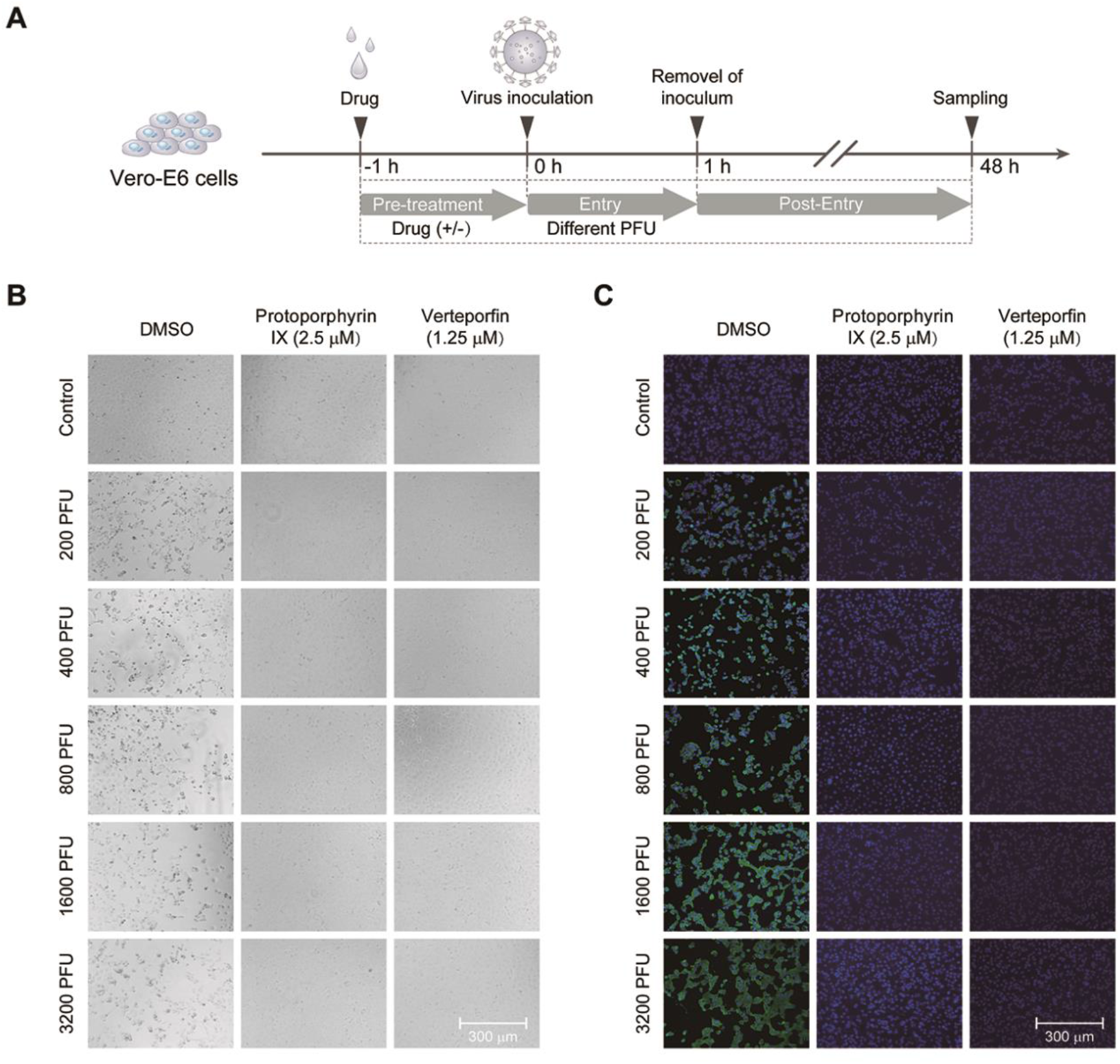
Protoporphyrin IX and verteporfin prevent SARS-CoV-2 infection. **(A)** Schematic presentation of treatment design. Briefly, Vero-E6 cells were pre-treated with protoporphyrin IX, verteporfin or the solvent DMSO before viral infection for 1 hour, then the drugs were removed and the cells were washed and infected with an increasing titer of SARS-CoV-2. **(B)** CPE of the cells with the different treatment. **(C)** Immunofluorescence of intracellular viral N protein. Intracellular expression of N protein of different treatment was assessed by staining of infected Vero-E6 cells with the polyclonal anti-N antibody (1:1000 dilution, green). Nuclei were stained with DAPI.

### Protoporphyrin IX and verteporfin interact with human ACE2 protein

Protoporphyrin IX and verteporfin share a same structure formed by four pyrrole rings (Fig. 4A) and thus likely act through a common antiviral mechanism. The above results suggest that both drugs act by inhibiting an early step in viral infection. One possibility was the saturation or modification of an essential cellular factor(s) required for viral infection. We thus investigated firstly by molecular docking analysis whether human ACE2, the viral receptor, might be the target of the compounds. The ACE2 peptidase domain (PD) from the human ACE2-B^0^AT1 complex (PDB ID: 6m18)^16^ was used for docking with protoporphyrin IX and verteporfin (Fig. 4A). The result with the highest ranking is exhibited in Fig. 4B, which represents the molecular model of protoporphyrin IX or verteporfin binding to PD. Protoporphyrin IX is located in the shallow-pocket-like space in the PD, with a binding energy of −5.60 kcal/mol. Similar result was obtained from the docking of verteporfin with PD (with a binding energy of −5.35 kcal/mol). Fig. 4C provides a view of the interaction of protoporphyrin IX or verteporfin with ACE2 PD residues. In the model, 25 residues of the PD interacted with protoporphyrin IX, in which the benzene ring of Phe^40^ interacted closely with the porphyrin-ring of protoporphyrin IX, the Trp^69^ formed aromatic H-bonds with the porphyrin-ring, Asp^350^ and Asp^382^ formed H-bonds with the compound. The other residues involved in the interaction with protoporphyrin IX included Ser^43^, Ser^44^, Ser^47^, Asn^51^, Gly^66^, Ser^70^, Leu^73^, Thr^347^, Ala^348^, Trp^349^, Leu^351^, Gly^352^, Phe^356^, His^378^, Ile^379^, Tyr^385^, Phe^390^, Leu^391^, Arg^393^, Asn^394^ and His^401^. Similar results were observed in the interaction between verteporfin and PD, except that Asn^51^ formed additional H-bonds with the benzazole-like structure of verteporfin. Many of these PD residues are located in the region that interacts with SARS-CoV-2 S protein receptor binding domain (RBD), especially Phe^40^, Ser^43^, Ser^44^, Trp^349^ - Gly^352^ and Phe^356^, which are very close to the key residues (Tyr^41^, Gln^42^, Lys^353^ and Arg^357^) that interact with the RBD^16^. The results suggest that protoporphyrin IX and verteporfin might interact with ACE2.

**Fig. 4.**
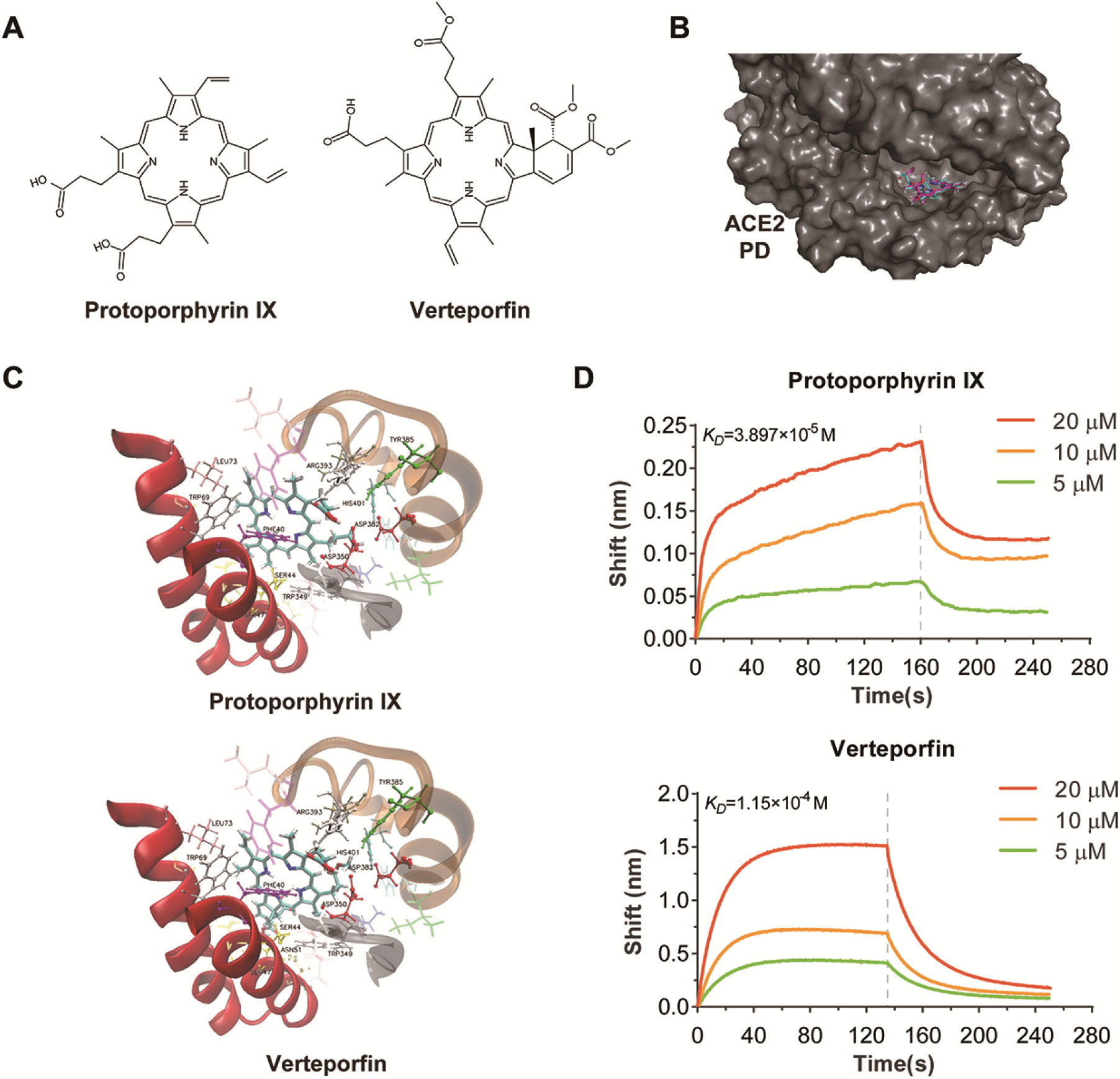
Protoporphyrin IX and verteporfin bind human ACE2 protein. **(A)** Structures of protoporphyrin IX and verteporfin. **(B)** Docking of ACE2 peptidase domain (PD) with protoporphyrin IX (blue) and verteporfin (pink). The 3D structure of PD is from cryo-electron microscopy structure of the ACE2-B^0^AT1 complex (PDB ID: 6m18). The surface of PD is shown. **(C)** Interactions of protoporphyrin IX (upper) or verteporfin (bottom) with ACE2 residues. **(D)** Binding profiles of protoporphyrin IX or verteporfin to ACE2-Fc protein measured with Biolayer Interferometry assay.

We next used BLI (Biolayer Interferometry) assay to evaluate the binding between ACE2 and these two compounds. As shown in Fig. 4D, protoporphyrin IX and verteporfin indeed bind to ACE2-Fc. The KD of protoporphyrin IX and verteporfin binding to ACE2-Fc were calculated to be 3.897×10^-5^ mol/L and 1.15×10^-4^ mol/L, respectively. Therefore, structural simulation by molecular docking and direct drug-protein binding assay support the binding of both drugs to viral receptor ACE2.

### Protoporphyrin IX and verteporfin interfere with the interaction between SARS-CoV-2 S protein and ACE2

Based on the molecular docking and the experimental data, both drugs likely interfere with the interaction between ACE2 and RBD via binding ACE2, which would impair viral entry. We first tested this possibility using a cell-cell fusion assay. HEK293T cells that express SARS-CoV-2 S protein served as the effector cells and those co-expressing human ACE2 and GFP as the target cells (Fig. 5A). The target cells were pre-treated with protoporphyrin IX (2.5 μM), verteporfin (1.25 μM) or DMSO for 1 hour. After removal of the drug, the target and effector cells were co-cultured at 37°C for 4 hours. Fused cells with larger cell size than normal cells were observed in the DMSO-treated group but barely in the protoporphyrin IX or verteporfin-treated group. The results indicate that protoporphyrin IX and verteporfin may block interaction of ACE2 and viral S protein which is required for cell-cell fusion.

**Fig. 5.**
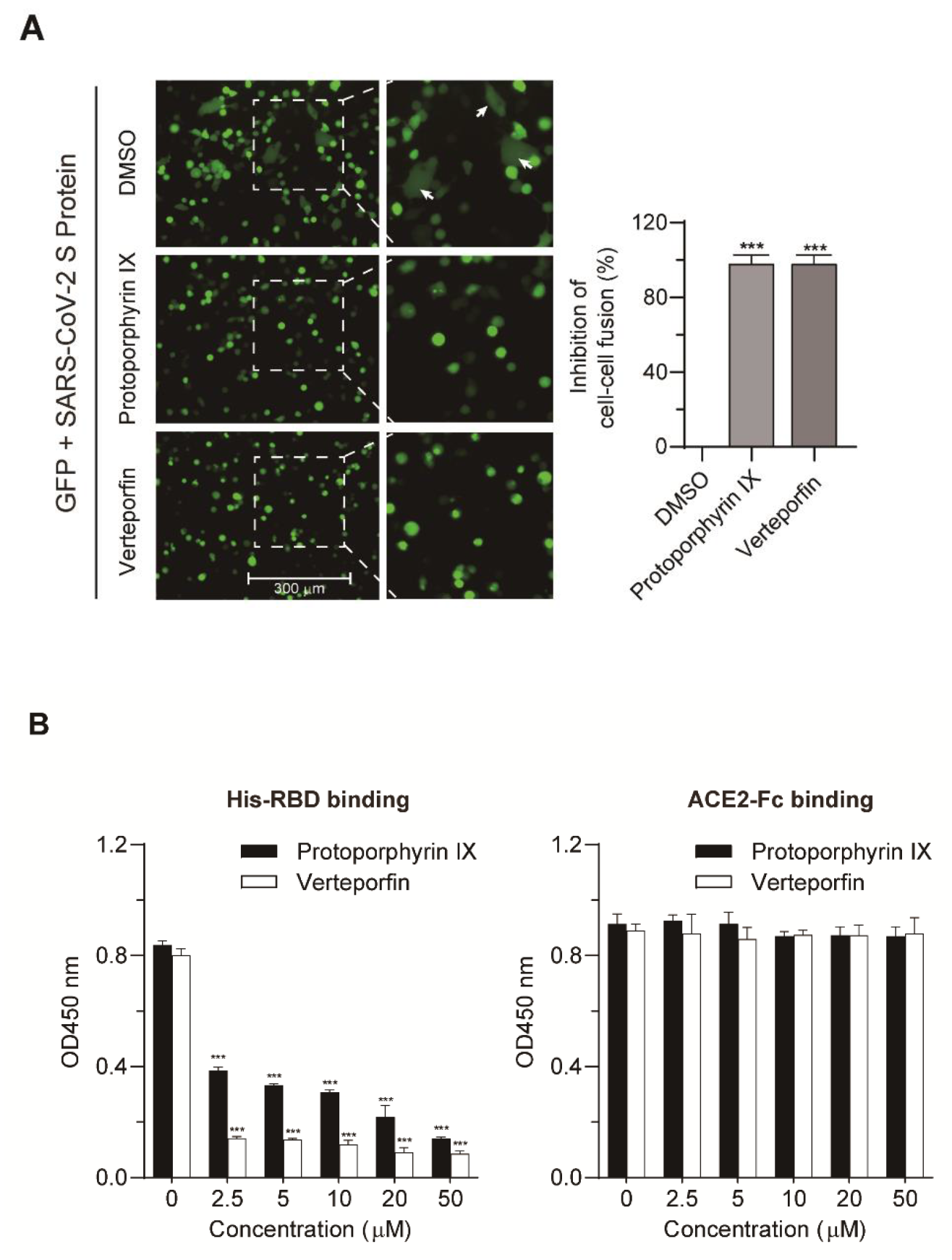
Protoporphyrin IX and verteporfin interfere with the interaction between ACE2 and RBD. **(A)** Blocking effect on ACE2 and SARS-CoV-2 S-mediated cell-cell fusion by protoporphyrin IX and verteporfin. The inhibitory value of protoporphyrin IX or verteporfin-treated group was presented relative to that of the DMSO-treated group which was set as 100%, respectively. Statistical significance was determined using the unpaired two-tailed Student’s t test. *** *P* < 0.001. **(B)** ELISA. The binding of His-RBD or ACE2-Fc to drug-treated pre-coated ACE-Fc or His-RBD was measured by absorbance at 450 nm. Statistical significance was determined using the unpaired twotailed Student’s *t* test. *** *P* < 0.001. Data from triplicate wells were analyzed.

To more directly demonstrate the interference of the compounds with the interaction of ACE2 to RBD, we designed an ELISA assay, in which protoporphyrin IX or verteporfin was added to 96-well plate pre-coated with ACE2-Fc or His-RBD. After incubation, unbound drugs were washed away. His-RBD or ACE2-Fc was added to the drug-treated wells pre-coated with ACE2-Fc or His-RBD. The results showed that both drugs could prevent the binding of His-RBD to pre-coated ACE2-Fc, while they had no effect on the binding of ACE2-Fc to pre-coated His-RBD (Fig. 5B). The data suggest that protoporphyrin IX and verteporfin most likely bind to ACE2 and interfere with the binding of RBD to ACE2, which is consistent with the results of the cell-cell fusion and molecular docking abovementioned.

### Protoporphyrin IX and verteporfin effectively prevent SARS-CoV-2 infection in the mouse model expressing human ACE2

To investigate the inhibition of protoporphyrin IX and verteporfin of SARS-CoV-2 infection *in vivo*, mice were first transduced intranasally with Ad5-hACE2 which could produce hACE2 in transduced HEK293T cells (Fig. S1). The mice were then infected intranasally with SARS-CoV-2 (2×10^5^ PFU/mouse) in a total volume of 50 μL DMEM containing protoporphyrin IX (100 μM), verteporfin (20 μM) or 1% DMSO (Fig. 6A). Incubation of protoporphyrin IX (100 μM) or verteporfin (20 μM) with the virus had no effect on viral infectivity (Fig. S2).

**Fig. 6.**
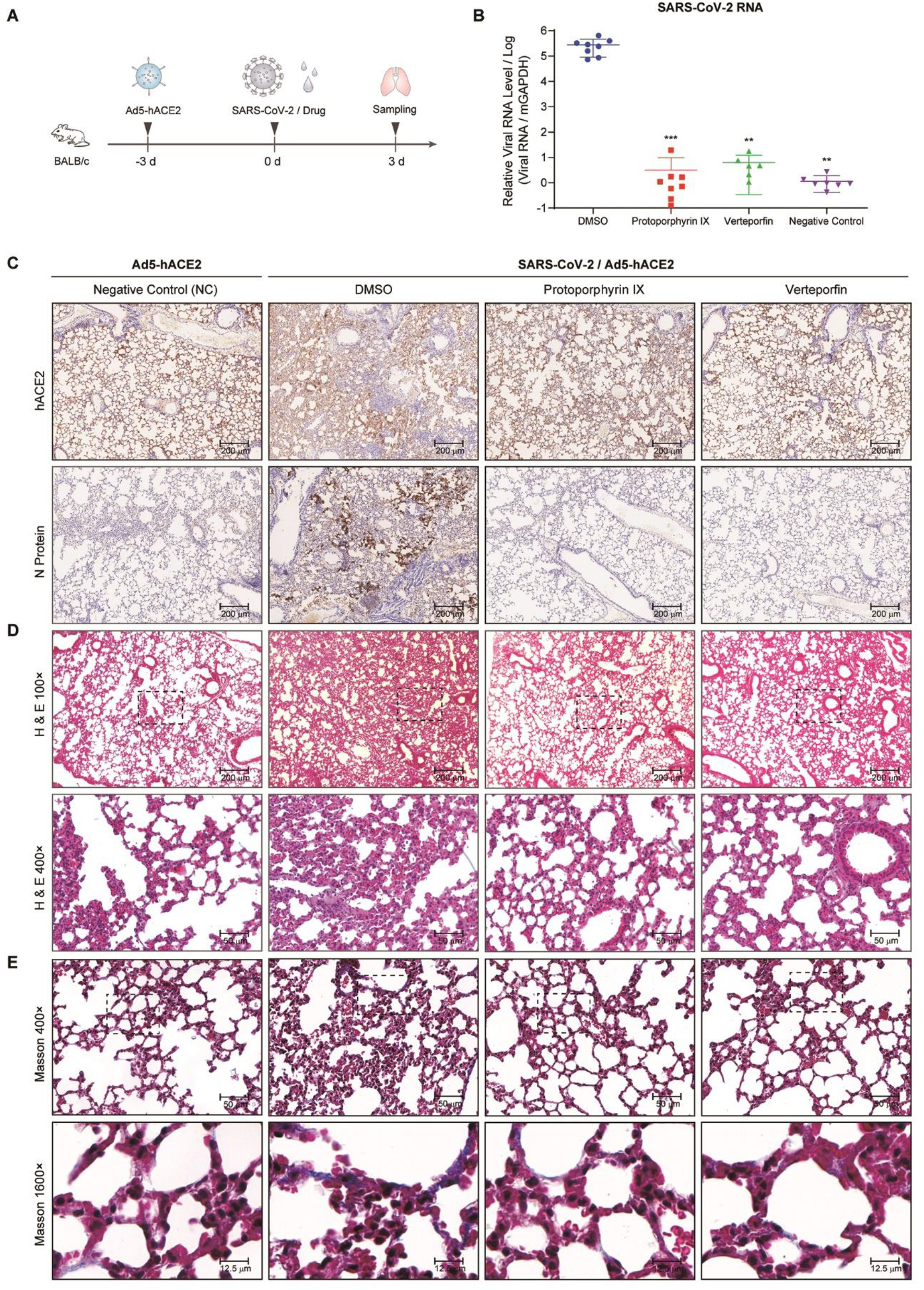
Effective inhibition of SARS-CoV-2 infection by protoporphyrin IX and verteporfin in SARS-CoV-2-infected hACE2 mice. **(A)** Schematic representation of the experiment timeline. **(B)** Relative viral RNA levels in lung tissues from each group. Data are relative to that of the DMSO-treated group and statistical significance was calculated using unpaired two-tailed t-test. \*\**P* < 0.01 and \*\*\**P* < 0.001. **(C)** Immunohistochemical staining of hACE2 and viral N protein in lung tissue samples from each group. **(D)** Representative HE staining of lung tissue sections from each group. **(E)** Representative Masson’s Trichrome staining of lung tissue sections from each group.

SARS-CoV-2-infected mice treated with 1% DMSO showed ruffled fur, hunching, loss of appetite and difficulty in breathing beginning 2 days post infection, while SARS-CoV-2-infected mice in the protoporphyrin IX and verteporfin groups were normal without obvious symptoms. All the mice were euthanized at day 3 post-infection of SARS-CoV-2 and lung tissues were collected. Human ACE2 expression in Ad5-hACE2 transduced mouse lung tissues was verified by immunochemical staining with the specific antibody, which lined along the pulmonary epithelial cells in DMSO group, protoporphyrin IX and verteporfin treated groups (Fig. 6C). Much fewer cells expressed viral N protein in the protoporphyrin IX and verteporfin groups compared to the DMSO group (Fig. 6C). Viral RNA was barely detected in most of the lung samples taken from the protoporphyrin IX and verteporfin groups and the mean of viral RNA levels from these groups were significantly lower than that from the DMSO group (Fig. 6B). The sections of lung tissues from the DMSO group displayed a variety of lesions including perivascular to interstitial inflammatory cell infiltrates, and necrotic cell debris. In contrast, the sections of lung tissues from the protoporphyrin IX and verteporfin groups showed no obvious histopathological change, so did those from the non-infected mice (the NC group) (Fig. 6D). The heavy deposition of collagen in the thickened alveolar interstitium was observed in the DMSO group with Masson-Trichrome staining, which was absent in the non-infected group and barely observed in the protoporphyrin IX and verteporfin groups (Fig. 6E). These results indicate that protoporphyrin IX and verteporfin also effectively inhibit SARS-CoV-2 infection in mouse model.

## Discussion

Protoporphyrin IX and verteporfin have been approved and used in the treatment of human diseases. Protoporphyrin IX is the final intermediate in the protoporphyrin IX iron complex (heme) biosynthetic pathway^17^. Heme is an important cofactor for oxygen transfer and oxygen storage^18^ and is a constituent of hemoproteins which play a variety of roles in cellular metabolism^19^. The light-activable photodynamic effect of protoporphyrin IX was used for cancer diagnosis^20^ and approved by FDA for treatment of bronchial and esophageal cancers and early malignant lesions of the skin, bladder, breast, stomach, and oral cavity^21,22^. Verteporfin was approved for the treatment of age-related macular degeneration^23^. The potential of verteporfin for the treatment of cancers, such as prostatic cancer, breast cancer, and pancreatic ductal adenocarcinoma has been investigated^24^. Verteporfin also has been reported to inhibit autophagy at an early stage by suppressing autophagosome formation^25^.

A study of clinical pharmacokinetics of verteporfin showed that in healthy volunteers who were infused with verteporfin 6 to 14 mg/m^2^ of body surface area over 1.5 to 45 minutes, Cmax (peak concentration) of verteporfin was 1.24-2.74 μg/ml^26^. The Cmax value is approximately 2.4 to 5.2-fold higher than the EC90 value that was obtained in this study (0.73 μM, i.e. 0.52 μg/ml). Protoporphyrin IX is the metabolite of 5-aminolevulinic acid (5-ALA) in human body. After administration of 5-ALA 2 mg/kg p.o., the average Cmax of protoporphyrin IX was 27.44 μg/ml^27^, which is about 20-fold higher than the EC90 value in this study (2.45 μM, i.e. 1.38 μg/ml). These data indicate that the two drugs can reach a plasma concentration that is much higher than the *in vitro* effective antiviral concentration. In the mouse model in this study, protoporphyrin IX and verteporfin exhibited effective inhibition of SARS-CoV-2 infection without notable toxicity.

Both protoporphyrin IX and verteporfin have a porphyrin ring structure formed by four pyrrole rings. It is most likely that they share a similar mechanism of antiviral action. In the experiment when either drug was added prior to viral infection, viral RNA production was inhibited even if the relevant drug was not added in the later virus infection and post-infection stages (group IV in Fig. 2B, 2C). Furthermore, increasing viral titer did not relieve the inhibition of the drugs added before viral infection (Fig. 3B). A logical hypothesis is that both drugs act by inhibiting an early step in viral infection. Structural simulation by molecular docking and direct drug-protein binding assay support the binding of both drugs to viral receptor ACE2. Several residues on ACE2 predicted to interact with the drugs are very close to the key residues that interact with the RBD of viral S protein. Based on the molecular docking and the experimental data, both drugs likely interfere with the interaction between ACE2 and RBD via binding ACE2, which would impair viral entry. The proposed mechanism was supported by the blocking effect of both drugs on the cell-cell fusion mediated by the interaction of ACE2 and viral S protein and by more direct evidence came from the ELISA binding assay. To our knowledge, this is the first report on small compounds that target the interaction between SARS-CoV-2 S protein and ACE2. The study suggests a new venue for the development of small molecule-based entry inhibitor against SARS-CoV-2. Furthermore, it may be a potential strategy for combating SARS-CoV-2 infections to use the compounds inhibiting virus entry in combination with the drugs acting intracellularly, such as the RdRp inhibitor remdesivir.

On the other hand, protoporphyrin IX and verteporfin were able to inhibit viral RNA production to some degree when they were added after viral infection (group VII in Fig. 2B, 2C). It is possible that the drugs might inhibit the infection of progeny viruses and hence prevent virus spreading. However, the absence of N protein expression in post-infection verteporfin-treated cells suggests that there might be other antiviral mechanism. Whether the drugs stimulate an antiviral innate immune response also needs exploration.

In conclusion, this study has discovered protoporphyrin IX and verteporfin as potent antiviral agents against SARS-CoV-2 infection in vitro and in the hACE2 mouse model. The effective antiviral concentrations of these drugs are in the nanomolar concentration range and the selectivity indices are greater than 200, indicating broad margin of safety. Both compounds bind viral receptor ACE2, thereby disturbing the interaction between ACE2 and the receptor-binding domain of viral S protein. To our knowledge, this is the first report on small compounds that target the interaction between SARS-CoV-2 S protein and ACE2, which sheds new light on developing novel chemoprophylaxis and chemotherapy against SARS-CoV-2. The antiviral efficacy of protoporphyrin IX and verteporfin *in vivo* will need clinical evaluation.

## Materials and Methods

### Cell line, virus, compounds and constructs

African green monkey kidney Vero-E6 cells and human embryonic kidney HEK293T cells were cultured at 37°C with 5% CO_2_ in Dulbecco’s modified Eagle medium (DMEM) (Gibco, Carlsbad, USA) containing 2 mmol/L L-glutamine, 50 U/mL penicillin, 100 mg/mL streptomycin, and 10% (vol/vol) fetal bovine serum (Gibco). Vero-E6 cells after SARS-CoV-2 infection were maintained in DMEM containing 2 mmol/L L-glutamine, 50 U/mL penicillin, 100 mg/mL streptomycin, and 2% (vol/vol) fetal bovine serum.

A clinical isolate of SARS-CoV-2, nCoV-SH01 (GenBank: MT121215.1)^28^, was propagated in Vero-E6 cells and the viral titer was determined as plaque forming units (PFU) per milliliter (mL) by CPE (cytopathic effect) quantification. All the infection experiments were performed in the biosafety level-3 (BSL-3) laboratory of Fudan University.

The recombinant adenovirus 5 expressing human ACE2 (Ad5-hACE2) and control adenovirus (Ad5-Ctrl) were purchased from ABM (Vancouver, Canada) or generated in the laboratory. For the generation of recombinant Ad5-hACE2, hACE2 cDNA was subcloned into the shuttle vector pShuttle-CMV^29^ between KpnI and XhoI sites, yielding pShuttle-CMV-hACE2. The plasmid pShuttle-CMV-hACE2 was linearized with restriction enzyme PmeI, and then transformed into BJ5183-AD-1 competent cells (Weidi, China), leading to the generation of pAd5-hACE2. Then, the plasmid pAd5-hACE2 was linearized with restriction enzyme PacI and used to transfect HEK293 cells as described previously^30^. Adenovirus Ad5-hACE2 was rescued from pAd5-hACE2-transfected cells and further amplified by several rounds of passage in HEK293 cells. High-titer adenovirus was purified by CsCl gradient centrifugation and virus titer was determined as described previously^31^. The resulting virus stock had a titer of 4.6*10^12^ vp/mL.

Custom compound libraries containing 3200 small molecules were purchased from Target Mol (MA, USA). Protoporphyrin IX (CAS No. 553-12-8), verteporfin (CAS No. 129497-78-5) and remdesivir (CAS No. 1809249-37-3) were purchased from MedChemExpress (NJ, USA).

pCMV-GFP and pcDNA3.1-ACE2 were constructed by inserting the green fluorescent protein (GFP) and human ACE2 cDNA into pcDNA3.1, respectively. pCAGGS-SARS-CoV-2-S that encodes the SARS-CoV-2 Spike gene was generated by GENEWIZ (Suzhou, China). Recombinant adenovirus expressing human ACE2 (Ad5-hACE2) and control adenovirus (Ad5-Ctrl) were purchased from ABM (Vancouver, Canada).

### Cell cytotoxicity assay

The Cell Counting Kit-8 (Dojindo, Kumamoto, Japan) was used to assess cell viability according to the manufacturer’s instructions. Briefly, Vero-E6 cells were dispensed into 96-well plate (1.0 x 10^4^ cells/well), cultured in medium supplemented with different concentrations of the compound for 48 hours. After removal of the medium, the cells were incubated with fresh serum-free medium containing 10% CCK-8 for 1 hour at 37°C and then the absorbances at 450 nm were measured using a microplate reader (Bio-Rad, Hercules, USA).

### Library screening

Custom compound libraries were screened via observation of CPE. Vero-E6 cells cultured in 96-well plate (4.0 x 10^4^ cells/well) were incubated with medium containing SARS-CoV-2 (200 PFU/well) and each compound (10 μM). Remdesivir (10 μM) served as positive control and DMSO as solvent control. CPE was observed under microscope every 24 hours for 72 hours.

### Evaluation of antiviral effects of the compounds

Vero-E6 cells cultured in 96-well plate (4.0 x 10^4^ cells/well) were pre-treated with the compound of a tested concentration or DMSO for 1 hour. SARS-CoV-2 (200 PFU/well) diluted in medium supplemented with the compound of the corresponding concentration was added and allow viral infection for 1 hour at 37°C. The mixture was removed and cells were washed twice with PBS, followed by culture with fresh medium containing the compound of the corresponding concentration. At 48 hours post infection, culture supernatant was collected for viral RNA quantification and the cells were fixed in 4% paraformaldehyde for immunofluorescence analysis.

To evaluate the relationship between the timing of compound addition and the antiviral efficacy, Vero-E6 cells cultured in 96-well plate (4.0 x 10^4^ cells/well) were treated with protoporphyrin IX (2.5 μM), verteporfin (1.25 μM) or DMSO at different timepoints relative to virus infection (Fig. 2a). Briefly, four sets of cells (I-IV) were pre-treated with the compound for 1 hour prior to virus infection. The medium was discarded and the cells were washed twice with PBS. Two sets (I, II) were then incubated with medium containing SARS-CoV-2 (200 PFU/well) and the compound for 1 hour and the other two sets (III, IV) were incubated only with the virus. After the removal of the virus and wash with PBS, set I and III were cultured with fresh medium containing the compound while set II and IV with medium without the compound. Four more sets of cells (V-VIII) were set up similarly except the initial medium contains DMSO instead of the compound. At 48 hours post infection, the culture supernatant was collected for viral RNA quantification and the cells for immunofluorescence analysis.

For evaluation of the prevention of viral infection by the compounds, Vero-E6 cells plated in 96-well plate (4.0 x 10^4^ cells/well) were pre-treated with protoporphyrin IX (2.5 μM), verteporfin (1.25 μM) or DMSO for 1 hour. The compound was removed and the cells were washed with PBS twice. Subsequently, the cells were incubated with medium containing an increasing dose of SARS-CoV-2 for 1 hour. After removal of the virus and wash with PBS, the cells were cultured for 48 hours for immunofluorescence analysis.

For evaluation of the possible inactivation of SARS-CoV-2 by the compounds, SARS-CoV-2 (2×10^5^ PFU) were treated with 1% DMSO, protoporphyrin IX (100 μM), verteporfin (20 μM) or 0.2% Triton X-100 for 30 minutes at room temperature. The compounds were removed through centrifugal ultrafiltration (30 kDa, Millipore, Darmstadt, Germany) and viral titers were measured with TCID50 assay on Vero-E6 cells.

### Viral RNA extraction and quantitative real time PCR (qRT-PCR)

Viral RNA in tissue and cell supernatant was extracted using TRIzol reagent (Invitrogen, Carlsbad, USA) following the manufacturer’s instructions. After phenol/chloroform extraction and isopropanol precipitation, RNA was reverse transcribed using cDNA Synthesis Kit (Tiangen, Shanghai, China) according to the manufacturer’s instructions. Quantitative real-time PCR (qRT-PCR) was performed in a 20 μL reaction containing SYBR Green (TaKaRa, Kusatsu, Japan) on MXP3000 cycler (Stratagene, La Jolla, USA) with the following program: initial denaturation at 95°C for 300 seconds; 40 cycles of 95°C for 15 seconds, 55°C for 20 seconds, and 72°C for 20 seconds; followed by a melt curve step. The PCR primers (Genewiz) targeting the N gene (nt608-706) of SARS-CoV-2 were: 5’-GGGGAACTTCTCCTGCTAGAAT-3’/5’-CAGACATTTTGCTCTCAAGCTG-3’ (forward/reverse).

### Immunofluorescence analysis

To detect the viral nucleocapsid protein (N protein), anti-N polyclonal antibodies were generated using standard immunization of BALB/c mice with recombinant N protein derived from *E. coli*. Vero-E6 cells grown in 96-well plate were fixed in 4% paraformaldehyde, permeabilized by 0.2% Triton X-100 (Thermo Fisher Scientific, Waltham, USA), blocked with 3% BSA, and stained overnight with the anti-N antibody (1:1000 dilution) at 4°C. The samples were then incubated with Alexa Fluor donkey anti-mouse IgG 488-labeled secondary antibody (1:1000 dilution, Thermo Fisher Scientific) for 1 hour at 37°C. The nuclei were stained with DAPI (Thermo Fisher Scientific). Images were captured with fluorescence microscopy (Thermo Fisher Scientific).

### Molecular docking

Cryo-electron microscopy structures of the full-length human ACE2 and a neutral amino acid transporter B^0^AT1 complex with an overall resolution of 2.9 Å have been reported^16^. The structure files were downloaded from Protein Data Bank (PDB ID: 6m18). Meanwhile, the structures of the compounds, protoporphyrin IX and verteporfin, were obtained from the EMBL-EBI and PubChem compound databases.

The receptor-ligand docking of the ACE2 protein with protoporphyrin IX or verteporfin was performed by using AutoDock 4.2.6 software and visualized with AutoDockTools 1.5.6 software (http://autodock.scripps.edu). Firstly, the ligand and receptor coordinate files were prepared respectively to include the information needed by AutoDock and the PDBQT files were created. Then the three-dimension of the grid box was set in AutoDockTools to create the grid parameter file. Afterwards, AutoGrid was used to generate the grid maps and AutoDock was run for receptor-ligand docking. After docking was completed, the results were shown in AutoDockTools, then the binding energy and receptor-ligand interactions were evaluated. The docking area was displayed in VMD 1.9.3 software (http://www.ks.uiuc.edu/Research/vmd).

### Cell-cell fusion assay

Cell-cell fusion was performed as described previously^32^. Briefly, target HEK293T cells were transiently co-transfected with pCMV-eGFP and pcDNA3.1-ACE2 using polyethyleneimine (PEI). Effector HEK293T cells were generated by transfection with the envelope-expressing plasmid pCAGGS-SARS-CoV-2-S. Twenty-four hours post transfection, the effector cells were pre-treated with protoporphyrin IX (2.5 μM), verteporfin (1.25 μM) or DMSO for 1 hour. The compound was then removed and the cells were washed with PBS twice. The target cells were quickly trypsinized and added to adherent effector cells in a 1:1 target-to-effector cell ratio. After a 4-hour co-cultivation period, five fields were randomly selected in each well and the number of fused and unfused cells in each field were counted directly under an inverted fluorescence microscope, based on much larger cell size of fused cells.

### ELISA

In the binding assay of viral S protein receptor binding domain (RBD), the recombinant protein of the extracellular domain of human ACE2 (aa 1-740) fused to Fc (ACE2-Fc, Genscript, Nanjing, China) was coated on 96-well microtiter plate (50 ng/well) at 4°C overnight. The wells were blocked with 3% BSA for 1 hour at 37°C. Serial dilution solutions of protoporphyrin IX, verteporfin or DMSO were added and incubated at 37°C for 1 hour. The free drug or DMSO was washed away with PBS. 50 ng of His-tagged RBD (His-RBD, aa 319-541) (Genscript) was then added to each well and incubated at 37°C for 2 hours. The wells were then washed with PBS and incubated with mouse anti-His antibody (1:1000 dilution, Abmart, Berkeley Heights, USA) at 37°C for 1 hour, followed by incubation with Horseradish peroxidase (HRP)-conjugated goat anti-mouse antibody (Abmart) at 37°C for 1 hour. Finally, TMB substrate was added for color development and the absorbance at 450 nm was read on a 96-well plate reader. The binding assay of ACE2 was similarly performed, except that His-RBD protein (50 ng/well) was coated on 96-well microtiter plate and ACE2-Fc protein was used for binding. HRP-goat anti-human Fc antibody (Abmart) was used for final signal detection.

### Transduction of HEK293T cells and Western blot analysis

HEK293T cells were transduced with Ad5-hACE2 or Ad5-Ctrl at a multiplicity of infection (MOI) = 100 for 4 hours at 37°C. The cells were lysed 48 hours post transduction and the samples were subjected to 10% SDS-PAGE and transferred to nitrocellulose membranes. The membranes were blocked with 3% bovine serum albumin (BSA) in PBST (PBS containing 0.05% Tween 20, pH7.0) and incubated with human ACE2 Rabbit Polyclonal antibody (1:100 dilution, Proteintech, Wuhan, China) followed by HRP-conjugated goat anti-rabbit IgG secondary antibody (1:5000 dilution, Invitrogen). Immobilon Western Chemiluminescent HRP Substrate (Thermo Fisher Scientific) was used for signal development.

### Transduction and infection of mice

Eight-week-old male mice (BALB/c) (SLAC Laboratory Animal, Shanghai, China) were raised in pathogen-free cages in the BSL-3 laboratory of Fudan University. The animal study protocol has been approved by the Animal Ethics Committee of School of Basic Medical Sciences, Fudan University.

Mice were transduced intranasally with Ad5-hACE2 (5×10^10^ viral particles per mouse in 50 μl saline) and were randomly divided into four groups three days post transduction. The mice were then infected intranasally with SARS-CoV-2 (2×10^5^ PFU per mouse) in a total volume of 50 μl DMEM containing 100 μM protoporphyrin IX (protoporphyrin IX group), 20 μM verteporfin (verteporfin group) or 1% DMSO (mock group), respectively. Non-SARS-CoV-2 infected Ad5-hACE2 transduced mice were used as negative control group (NC group). Mice were monitored and weighed daily. All the mice were euthanized and sacrificed at day 3 post infection to collect the lungs for the examinations of virus infection and histopathological changes.

### Preparation of lung tissue samples

Mouse lung tissues were fixed in 4% paraformaldehyde solution. Tissue homogenates (1 g/mL) were prepared by homogenizing perfused lung tissues using an automatic sample grinding instrument (Jingxin, Shanghai, China) for 1 minute in TRIzol reagent. The homogenates were centrifuged at 12,000 rpm for 10 minutes at 4 °C. The supernatant was collected for viral RNA extraction.

### Histology and immunohistochemistry

Mouse lungs were fixed in 4% paraformaldehyde solution. Tissue paraffin sections (2~4 μm in thickness) were stained with hematoxylin and eosin (H&E) and modified Masson’s Trichrome. To detect hACE2 expression, the sections were first incubated in blocking reagent and then with hACE2 Rabbit Polyclonal antibody (1:100 dilution, Proteintech) at 4 °C overnight, followed by incubation with HRP-conjugated goat anti-rabbit IgG secondary antibody (1:5000 dilution, Invitrogen). The lung sections from the mouse transduced intranasally with 5×10^10^ of Ad5-hACE2 were used as negative control. For viral antigen detection, the sections were sequentially incubated with mouse polyclonal antibody to SARS-CoV-2 N protein (1:500 dilution) and HRP-conjugated goat anti-mouse IgG secondary antibody (1:5000 dilution, Invitrogen). The sections were observed under microscope (Olympus, Tokyo, Japan).

### Biolayer interferometry (BLI) binding Assay

BLI assays were carried out in 96-well black plates using an OctetRED96 device (Pall ForteBio, Fremont, USA). For detecting the binding kinetics of protoporphyrin IX or verteporfin with hACE2, the recombinant protein ACE2-Fc (Genscript) at 5 μg/mL buffered in PBST (PBS with 0.02% Tween 20, pH 7.0) was immobilized onto activated AHC biosensors (ForteBio) and incubated with 20 μM, 10 μM or 5 μM of each compound in kinetics buffer (PBST). The experiment included the following steps at 37°C: (1) equilibration (60 seconds); (2) immobilization of ACE2-Fc onto sensors (100 seconds); (3) baseline in kinetics buffer (60 seconds); (4) association of the drug for measurement of kon (240 seconds); and (5) dissociation of the drug for measurement of koff (200 seconds). All the curves were fitted by a 2:1 (heterogeneous ligands) binding model and mean K_D_ values were determined using the Data Analysis software (ForteBio).

### Statistical analysis

Data were analyzed using Prism 7 (GraphPad) and were presented as mean ± SEM. The dose response curves of viral RNA levels or cell viability *versus* the drug concentrations were plotted and evaluated by Prism 7. Statistical significance was determined using unpaired two-tailed Student’s *t* test for single variables and two-way ANOVA followed by Bonferroni posttests for multiple variables.

## Acknowledgments

The study was supported by the National Science and Technology Major Project (NSTMP) for the Prevention and Treatment of Infectious Diseases (2018ZX10734401, 2018ZX10301208), NSTMP for the Development of Novel Drugs (2019ZX09721001), and Project of Novel Coronavirus Research of Fudan University.

## Author Contributions

Youhua Xie, Di Qu and Qing Deng drafted the manuscript. Youhua Xie, Di Qu and Qiang Deng designed the project. The majority of the experiments and data analysis were performed by Chenjian Gu, Yang Wu, Huimin Guo and Yuanfei Zhu. The other authors participated in the data analysis and manuscript revision. All the authors have approved the manuscript.

---

**Figure S1.**
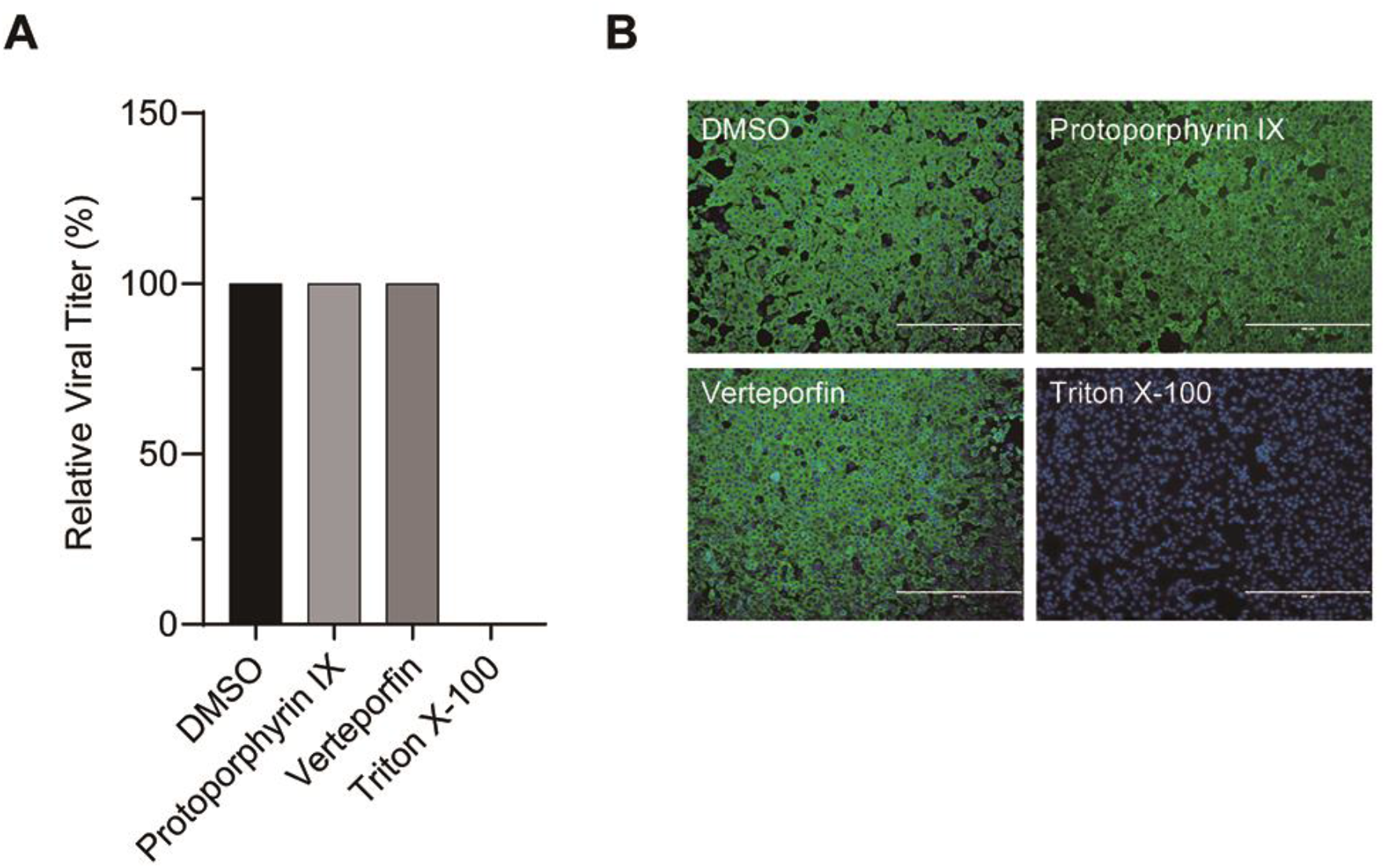
Incubation of protoporphyrin IX or verteporfin with SARS-CoV-2 has no effect on viral infectivity. **(A)** Relative viral titer measured with TCID50 assay. SARS-CoV-2 (2×10^5^ PFU) was treated with 1% DMSO, protoporphyrin IX (100 μM), verteporfin (20 μM) or 0.2% Triton X-100 for 30 minutes. The compounds were removed by centrifugal ultrafiltration and viral titers were measured with TCID50 assay on Vero-E6 cells. (**B**) Immunofluorescence of intracellular viral N protein. Intracellular expression of N protein was assessed by staining infected Vero-E6 cells using the polyclonal anti-N antibody (1:1000 dilution, green). Nuclei were stained with DAPI.

**Figure S2.**
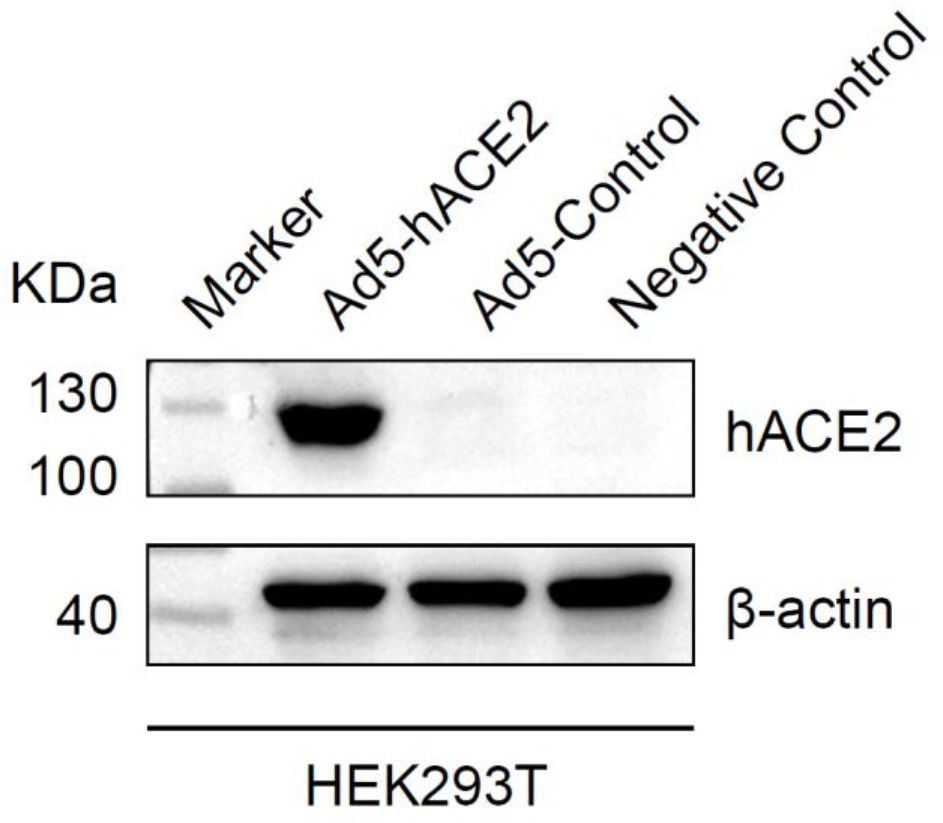
Expression of hACE2 in Ad5-hACE2 transduced HEK293T cells. Western blot analysis of hACE2 protein in HEK293T cells transduced by Ad5-hACE2. HEK293T cells were transduced with Ad5-hACE2 or Ad5-Ctrl at MOI of 100 at 37 °C for 4 hours. HEK293T cells without any treatment were used as negative control.

## References

1 Elfiky, A. A. Ribavirin, Remdesivir, Sofosbuvir, Galidesivir, and Tenofovir against SARS-CoV-2RNA dependent RNA polymerase (RdRp): A molecular docking study. Life Sciences 253, 117592, doi:https://doi.org/10.1016/j.lfs.2020.117592 (2020).

2 Wang, M. et al. Remdesivir and chloroquine effectively inhibit the recently emerged novel coronavirus (2019-nCoV) in vitro. Cell Res 30, 269–271, doi:10.1038/s41422-020-0282-0 (2020).

3 Grein, J. et al. Compassionate Use of Remdesivir for Patients with Severe Covid-19. New England Journal of Medicine, doi:10.1056/NEJMoa2007016 (2020).

4 Wang, Y et al. Remdesivir in adults with severe COVID-19: a randomised, double-blind, placebo-controlled, multicentre trial. Lancet 395, 1569–1578, doi:10.1016/S0140-6736(20)31022-9 (2020).

5 Beigel, J. H. et al. Remdesivir for the Treatment of Covid-19 - Preliminary Report. N Engl J Med, doi:10.1056/NEJMoa2007764 (2020).

6 Costanzo, M., De Giglio, M. A. R. & Roviello, G. N. SARS-CoV-2: Recent Reports on Antiviral Therapies Based on Lopinavir/Ritonavir, Darunavir/Umifenovir, Hydroxychloroquine, Remdesivir, Favipiravir and Other Drugs for the Treatment of the New Coronavirus. Curr Med Chem, doi:10.2174/0929867327666200416131117 (2020).

7 Tu, Y F. et al. A Review of SARS-CoV-2 and the Ongoing Clinical Trials. Int J Mol Sci 21, doi:10.3390/ijms21072657 (2020).

8 Caly, L., Druce, J. D., Catton, M. G., Jans, D. A. & Wagstaff, K. M. The FDA-approved drug ivermectin inhibits the replication of SARS-CoV-2 in vitro. Antiviral Res 178, 104787 (2020).

9 Gautret, P. et al. Hydroxychloroquine and azithromycin as a treatment of COVID-19: results of an open-label non-randomized clinical trial. Int J Antimicrob Agents, 105949 (2020).

10 Cortegiani, A., Ingoglia, G., Ippolito, M., Giarratano, A. & Einav, S. A systematic review on the efficacy and safety of chloroquine for the treatment of COVID-19. J Crit Care, doi:10.1016/j.jcrc.2020.03.005 (2020).

11 Suranagi, U. D., Rehan, H. S. & Goyal, N. Hydroxychloroquine for the management of COVID-19: Hope or Hype? A Systematic review of the current evidence. medRxiv, 2020.2004.2016.20068205, doi:10.1101/2020.04.16.20068205 (2020).

12 Magagnoli, J. et al. Outcomes of hydroxychloroquine usage in United States veterans hospitalized with Covid-19. medRxiv, 2020.2004.2016.20065920, doi:10.1101/2020.04.16.20065920 (2020).

13 Alsoussi, W. B. et al. A Potently Neutralizing Antibody Protects Mice against SARS-CoV-2 Infection. J Immunol, doi:10.4049/jimmunol.2000583 (2020).

14 Chi, X. et al. A neutralizing human antibody binds to the N-terminal domain of the Spike protein of SARS-CoV-2. Science, doi:10.1126/science.abc6952 (2020).

15 Shi, R. et al. A human neutralizing antibody targets the receptor-binding site of SARS-CoV-2. Nature, doi:10.1038/s41586-020-2381-y (2020).

16 Yan, R. et al. Structural basis for the recognition of SARS-CoV-2 by full-length human ACE2. Science 367, 1444–1448, doi:10.1126/science.abb2762 (2020).

17 Sachar, M., Anderson, K. E. & Ma, X. Protoporphyrin IX: the Good, the Bad, and the Ugly. J Pharmacol Exp Ther 356, 267–275 (2016).

18 Shimizu, T., Lengalova, A., Martinek, V. & Martinkova, M. Heme: emergent roles of heme in signal transduction, functional regulation and as catalytic centres. Chem Soc Rev 48, 5624–5657, doi:10.1039/c9cs00268e (2019).

19 Smith, L. J., Kahraman, A. & Thornton, J. M. Heme proteins--diversity in structural characteristics, function, and folding. Proteins 78, 2349–2368, doi:10.1002/prot.22747 (2010).

20 Ishizuka, M. et al. Novel development of 5-aminolevurinic acid (ALA) in cancer diagnoses and therapy. Int Immunopharmacol 11, 358–365, doi:10.1016/j.intimp.2010.11.029 (2011).

21 Pass, H. I. Photodynamic therapy in oncology: mechanisms and clinical use. J Natl Cancer Inst 85, 443–456, doi:10.1093/jnci/85.6.443 (1993).

22 Oleinick, N. L. & Evans, H. H. The photobiology of photodynamic therapy: cellular targets and mechanisms. Radiat Res 150, S146–156 (1998).

23 Schmidt-Erfurth, U. & Hasan, T. Mechanisms of action of photodynamic therapy with verteporfin for the treatment of age-related macular degeneration. Surv Ophthalmol 45, 195–214, doi:10.1016/s0039-6257(00)00158-2 (2000).

24 Pellosi, D. S. et al. Multifunctional theranostic Pluronic mixed micelles improve targeted photoactivity of Verteporfin in cancer cells. Mater Sci Eng C Mater Biol Appl 71, 1–9, doi:10.1016/j.msec.2016.09.064 (2017).

25 Donohue, E. et al. Inhibition of autophagosome formation by the benzoporphyrin derivative verteporfin. J Biol Chem 286, 7290–7300 (2011).

26 Houle, J. M. & Strong, A. Clinical pharmacokinetics of verteporfin. J Clin Pharmacol 42, 547–557, doi:10.1177/00912700222011607 (2002).

27 Leaders, T. & Uppoor, R. Clinical Pharmacology and Biopharmaceutics Review.

28 Rong, Z. et al. Isolation of a 2019 novel coronavirus strain from a coronavirus disease 19 patient in Shanghai. JOURNAL OF MICROBES AND INFECTIONS 15, 111–121 (2020).

29 Luo, J. et al. A protocol for rapid generation of recombinant adenoviruses using the AdEasy system. Nat Protoc 2, 1236–1247, doi:10.1038/nprot.2007.135 (2007).

30 Chi, Y. D. et al. Survivin-targeting Artificial MicroRNAs Mediated by Adenovirus Suppress Tumor Activity in Cancer Cells and Xenograft Models. Mol Ther-Nucl Acids 3, doi:ARTN e20810.1038/mtna.2014.59 (2014).

31 Li, G. et al. Recombinant covalently closed circular DNA of hepatitis B virus induces long-term viral persistence with chronic hepatitis in a mouse model. Hepatology 67, 56–70, doi:10.1002/hep.29406 (2018).

32. Xia, S. et al. Inhibition of SARS-CoV-2 (previously 2019-nCoV) infection by a highly potent pan-coronavirus fusion inhibitor targeting its spike protein that harbors a high capacity to mediate membrane fusion. Cell Res 30, 343–355, doi:10.1038/s41422-020-0305-x (2020).

